# Interacting phenotypes and the coevolutionary process: Interspecific indirect genetic effects alter coevolutionary dynamics

**DOI:** 10.1101/2021.04.14.439826

**Authors:** Stephen P. De Lisle, Daniel I. Bolnick, Edmund D. Brodie, Allen J. Moore, Joel W. McGlothlin

**Affiliations:** Department of Ecology & Evolutionary Biology, University of Connecticut, 75 N. Eagleville Road, Storrs, Connecticut, USA 06269; Evolutionary Ecology Unit, Department of Biology, Lund University, Solvegatan 37, Lund, Sweden; Department of Biology and Mountain Lake Biological Station, University of Virginia, 485 McCormick Road, Charlottesville, VA 22904 USA; Department of Entomology, University of Georgia, Athens, GA 30602 USA; Department of Biological Sciences, Virginia Tech, 2125 Derring Hall, 926 West Campus Drive, Blacksburg, Virginia, USA 24060

**Keywords:** Coevolution, cross-species selection, interspecific indirect genetic effects, quantitative genetics, species interactions

## Abstract

Coevolution occurs when species interact to influence one another’s fitness, resulting in reciprocal evolutionary change. In many coevolving lineages, trait expression in one species is modified by the genotypes and phenotypes of the other, forming feedback loops reminiscent of models of intraspecific social evolution. Here, we adapt the theory of within-species social evolution, characterized by indirect genetic effects and social selection imposed by interacting individuals, to the case of interspecific interactions. In a trait-based model, we derive general expressions for multivariate evolutionary change in two species and the expected between-species covariance in evolutionary change when selection varies across space. We show that reciprocal interspecific indirect genetic effects can dominate the coevolutionary process and drive patterns of correlated evolution beyond what is expected from direct selection alone. In extreme cases, interspecific indirect genetic effects can lead to coevolution when selection does not covary between species or even when one species lacks genetic variance. Moreover, our model indicates that interspecific indirect genetic effects may interact in complex ways with cross-species selection to determine the course of coevolution. Importantly, our model makes empirically testable predictions for how different forms of reciprocal interactions contribute to the coevolutionary process.

## Introduction

Coevolution occurs when interacting lineages evolve reciprocally in response to one another (Janzen 1980, Thompson 1982). Although the concept of coevolution may be applied to lineages that share genes, such as males and females of the same species (Arnqvist and Rowe 2002), it was originally invoked to explain patterns of correlated evolution between interacting species (Ehrlich and Raven 1964). In some cases, coevolution can result in tightly integrated mutualisms or spectacular arms races that drive the evolution of exceptional phenotypes (Brodie et al. 2002, Pellmyr 2003, Johnson and Anderson 2010). Yet even beyond these striking cases, coevolution is likely important for a wide range of interacting lineages, including consumers and their resources, hosts and their pathogens, competitors, and mutualists (Thompson 1982, 1994). Although coevolution has clearly played a major role in the origins of diversity, much is still unknown about when and how species interactions generate reciprocal evolutionary change.

Theoretical models of coevolution typically focus on the fitness effects of trait interactions between coevolving species and how selection imposed by one species manifests evolutionary change in an interacting species (Nuismer 2017). The interaction between species in coevolving lineages is often intimate, with one species spending a greater part of its life cycle in close contact with the other. Thus, coevolution bears a striking resemblance to intraspecific social evolution (Stearns 2012). Like social evolution, coevolution often includes interacting or extended phenotypes, which arise when traits can only be understood within the context of interactions with others (Dawkins 1982, Moore et al. 1997).

In quantitative genetic models of intraspecific social evolution, social interactants influence one another via two pathways, each of which has a counterpart in coevolutionary theory. First, the phenotype of one individual may cause fitness effects in a social partner, leading to a form of selection known as social selection (West-Eberhard 1979, West-Eberhard 1983, West-Eberhard 1984, Wolf et al. 1999). At the heart of all coevolutionary models is a form of reciprocal fitness interaction that resembles social selection, where the fitness of individuals in one species is influenced by traits in an interacting species (Brodie and Ridenhour 2003, Ridenhour 2005, Nuismer 2017). Second, models of social evolution may also include indirect genetic effects, which occur when the phenotype of one individual depends on the genotype of an interacting partner (Moore et al. 1997, Wolf et al. 1998). A classic example of indirect genetic effects is maternal effects, in which offspring phenotype is a function of both their own genes (a direct genetic effect) and maternal phenotypes such as litter size and provisioning (an indirect genetic effect) (Kirkpatrick and Lande 1989, Mousseau and Fox 1998, McAdam et al. 2002). When indirect genetic effects are reciprocal, feedback effects may inflate the genetic variance available for response to selection, drastically accelerating the rate of evolution (Moore et al. 1997). Although most models of indirect genetic effects do not extend beyond species boundaries, indirect genetic effects may also be common in species interactions between mutualists, competitors, parasites and hosts, and predators and prey. Phenotypic plasticity in response to an interacting species is common (Agrawal 2001), and when these influences on trait expression have a genetic basis they may represent interspecific indirect genetic effects (IIGEs; Shuster et al. 2006). Although IIGEs have received some attention in the context of community genetics (Shuster et al. 2006, Witham et al. 2020), their potential role in driving trait coevolution has been mostly unexplored.

To date, most explorations of IIGEs have been studies providing empirical support for their likely existence and their contribution to trait variation. Examples of interspecific phenotypic manipulation are common in nature (Table 1), and many of these cases can be argued to be putative cases of IIGEs. One possible example occurs in arbuscular mycorrhizae, where the genotypes of fungal mutualists can alter root traits in the plants they inhabit (Gianinazzi-Pearson et al. 2007). In host-parasite systems, parasite manipulation of host traits (such as behavior) and reciprocal host manipulation of parasite traits (such as growth rate, via immune response) are key features of species interactions (Thomas et al. 2012). Importantly, in many host-parasite systems, both the host and the parasite experience sustained interactions with a small number of individuals of the other species, often over key periods of the life history. For example, helminth parasites excrete a variety of immunomodulatory products that suppress or misdirect the immune system of their individual host (Damian 1997, Schmid-Hempel 2008, Oladiran and Belosevic 2012). Thus, host immune response to infection is controlled by the genotype of both the host and parasite (e.g., Barribeau 2014). As a specific example, some populations of threespine stickleback (*Gasterosteus aculeatus*) initiate a strong immune response to infection by the cestode *Schistocephalus solidus* (Fig. 1A) involving granulocyte proliferation and fibrosis, which effectively suppress cestode growth and viability (Weber et al. 2017). Other stickleback populations do not exhibit this response and allow rapid cestode growth, perhaps representing a tolerance strategy. Cestode growth is thus an indirect genetic effect of its host’s genotype. Conversely, the cestode has been shown to secrete compounds that suppress this host response (Scharsack et al. 2004, 2007, 2013) and down-regulate sticklebacks’ pro-fibrotic gene expression (Fuess et al. 2020), suggesting reciprocal indirect genetic effects.

**Figure 1.**
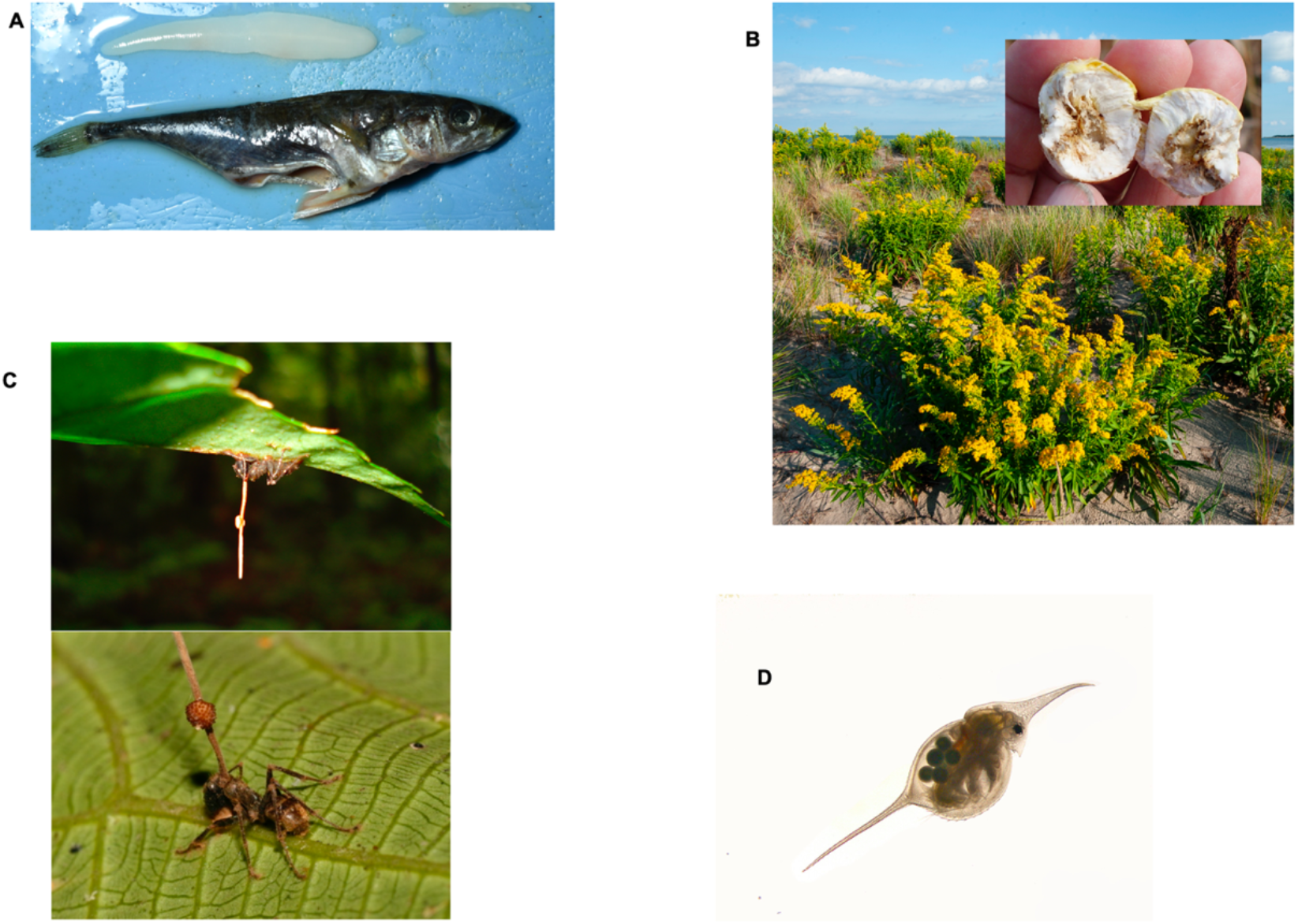
Examples of traits potentially mediated by IIGEs. IIGEs are a common feature of species interactions, particularly in host-parasite systems where prolonged contact between single individuals of both species determines fitness and trait expression for both. Here we highlight four examples where such IIGEs are likely contributing to trait expression and possibly patterns of among-population (co)variation. Panel A shows threespine stickleback fish (*Gasterosteus aculeatus*) and their cestode parasite *Schistocephalus*. Freshwater stickleback are the intermediate host for *Schistocephalus*, and each cestode acquires most of its lifetime resource pool while living inside a single host fish. *Schistocephalus* impose substantial reproductive and survival costs on hosts, and hosts have evolved an inducible (by the cestode) defense that suppresses cestode growth. Substantial among population-variation in this trait and in infection rates suggest these IIGEs may be mediating coevolution. Panel B shows goldenrod (*Solidago*) and a gall induced by the larvae (visible inside the gall) of the specialist gall-forming fly *Eurosta*. Gall size is induced by the genotype of the *Eurosta* larvae, and past work has shown that gall expression is a complex interaction between plant and fly genotype. Panel C shows the fungus *Ophiocordyceps unilateralis*, which manipulates behavior of its ant host prior to emergence of its fruiting body (Anderson et al. 2009). Panel D shows the water flea *Daphnia lumholtzi,* which induces growth of protective spines in response to chemical cues released by predatory fish (Agrawal 2001). Panel A and B main photos: S. De Lisle. Inset (B) photo: SriMesh / CC BY-SA (https://creativecommons.org/licenses/by-sa/3.0); C photo: “File:Ophiocordyceps unilateralis.png” by David P. Hughes, Maj-Britt Pontoppidan / CC BY 2.5; Panel D: “Water flea (Daphnia lumholtzi)” by Frupus / CC BY-NC 2.0.

**Table 1.** Coevolutionary interactions where IIGEs may be prevalent.

| Interaction | IIGE synonym | Diffuse/individual interaction | Example References |
| --- | --- | --- | --- |
| host-parasite | host manipulation | both | (Thomas et al. 2012) |
| predator-prey | trait-mediated indirect interaction | both | (Peacor and Werner 2001) |
| plant-animal | inducible direct defense<br>host-plant manipulations | both | (Weis and Abrahamson 1986, Chen 2008) |
| plant-microbe | joint trait<br>microbially mediated trait | diffuse | (Friesen et al. 2011, O'Brien et al. 2021) |
| host-microbiome | host control<br>host-microbiome interaction | diffuse | (Stappenbeck and Virgin 2016, Foster et al. 2017) |
| consumer-resource | toxin sequestration | both | (Züst et al. 2018) |

In goldenrod (*Solidago*; Fig. 1B), size of galls produced by the gall fly *Eurosta* is determined by genotypes of both fly and plant, and evolution of gall size is influenced in part by cross-species selection imposed on *Eurosta* larvae by species at other trophic levels (Weis and Abrahamson 1986, Weis et al. 1992, Abrahamson and Weis 1997). This type of interaction is common across gall-forming insects and their plant hosts; in *Hormaphis* aphids, variation the *bicycle* gene has been linked to variation in gall size (Korgaonkar et al. 2021). In many herbivore-plant interactions, physical damage to leaves induces upregulation of defensive compounds to deter further herbivory, which can be countered by matching physiological changes in the herbivore (Ohgushi 2005). For example, *Littorina* snail herbivory changes foliar chemistry of the brown seaweed *Ascophyllum nodosa* (increased phlorotannin concentrations), which in turn reduces snail movement and consumption rates (Borell et al. 2004).

Host-parasite, host-parasitoid, and some plant-herbivore interactions can entail intimate long-term associations between individuals. In contrast, predator-prey interactions tend to be more diffuse (Brodie and Brodie 1999). Prey sense predation risk through chemical, auditory, or visual cues and change their morphology, physiology, or behavior in ways that mitigate their risk of predator encounter (Werner and Peacor 2003, Preisser et al. 2005). Perhaps the most prominent example is the tendency of some *Daphnia* genotypes to grow spines (Fig. 1D), or to migrate to other depths, when they detect scent cues (kairomones) from predatory fish (Weber and Declerck 1997, Boersma et al. 1998). These antipredator responses lead to systemic changes in gene expression and morphology (Tams et al. 2019), which are controlled in part by fish traits (e.g., production of a scent cue). These scents can themselves be variable and genetic, as illustrated by differences in how *Daphnia* respond to cues from landlocked versus anadromous alewife (Walsh and Post 2011). The *Daphnia* traits can thus be described resulting from IIGEs controlled by both *Daphnia* and fish genotypes. Unlike the intimate host-parasite interactions, however, prey may be responding to diffuse cues from a predator population as a whole.

Although IIGEs would appear to play a central role in trait interactions between many coevolving species, little is known about how these effects influence the dynamics of coevolution (Scheiner et al. 2015). Past approaches, which have included variance-partitioning models of community assembly (Shuster et al. 2006, Whitham et al. 2020) and models of “joint traits” expressed together by interacting species (Queller 2014, O’Brien et al. 2021), are suggestive of an important role for IIGEs in species interactions. However, we lack a general understanding of how reciprocal IIGEs may affect the coevolution of interacting phenotypes. The fact that indirect genetic effects within a species can create feedback loops and other complex evolutionary dynamics, including in the context of within-species coevolution (Drown and Wade 2014), suggests that IIGEs could play a major role in mediating trait coevolution between species. Coopting concepts from intraspecific social evolution theory, where trait-based IGEs are well developed, thus provides a natural way to understand the trait interactions that drive coevolution. Importantly, this trait-based approach allows the contribution of indirect genetic effects to the coevolutionary process to be fully explored. Our goal in this paper is to develop such models to provide a comprehensive theoretical assessment of how IIGEs contribute to coevolution between species.

Here we adapt the trait-based theory of intraspecific social evolution (Moore et al. 1997, Wolf et al. 1999, McGlothlin et al. 2010) to the case of two interacting species. Our model applies to both pairwise interactions between individuals (e.g., stickleback and cestodes) as well as diffuse interactions between species mean values (e.g., predatory fish and *Daphnia*). Importantly, our model accommodates both IIGEs, where genes in individuals of one species influence trait expression in individuals of another species, and cross-species selection, where phenotypes of individuals in one species influence fitness of individuals of another species (Figs. 1, 2). In addition to describing the contribution of these interspecific interactions to evolutionary change, we develop expressions for the among-population evolutionary covariance between traits of two interacting species; that is, the expected covariance in trait means across populations between two coevolving species. In the process, we formalize an interspecific analog of Zeng’s (1988) quantitative genetic model of among-population trait covariation, which we expand to incorporate interacting phenotypes. Our analysis shows that IIGEs may have a central role in driving and mediating coevolution.

**Figure 2.**
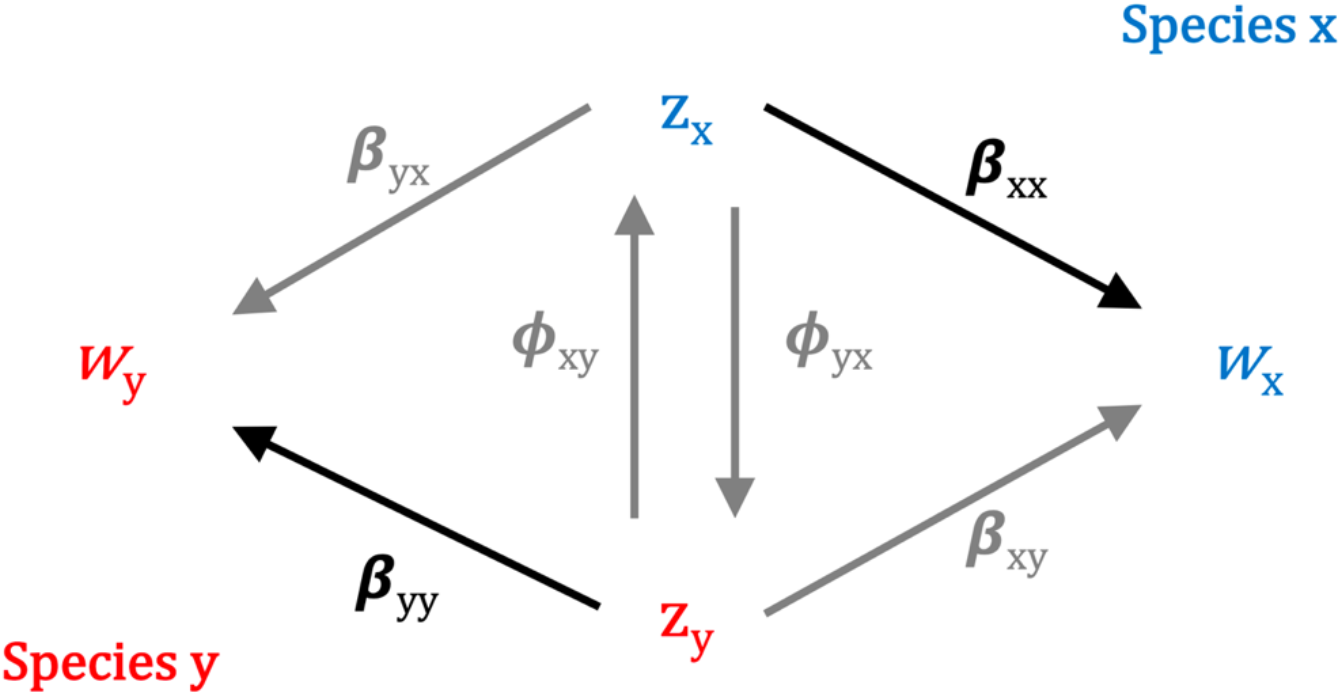
Path model of selection when traits and fitness are determined by interactions with heterospecifics. Individual trait values of two interacting species, *x* (blue) and *y* (red), are represented by *z*. Traits directly influence individual relative fitness (*w*) of the species that express them via natural selection (β_*xx*_ and β_*yy*_, black arrows). Traits can also reciprocally influence expression of traits of heterospecific social partners, via interspecific indirect genetic effects Φ. Traits can also influence fitness of heterospecific social partners, via cross-species selection (β_*xy*_ and β_*yx*_, grey arrows). Social effects are illustrated in grey arrows, direct effects in black.

### Reciprocal evolutionary change in interacting species

To model coevolution in two interacting species, we first decompose trait expression into three components: direct genetic effects, environmental effects, and indirect effects mediated by the phenotype of an interacting species. The phenotypic interface of coevolution involves traits with interacting effects across individuals of two species *x* and *y* and can be written as

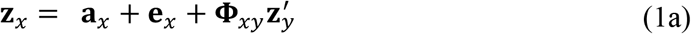

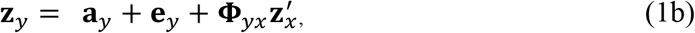

where **z**_*i*_ is a column vector of *m* traits expressed in an individual of species *i* = *x* or *y* (*m* is not necessarily equal in each species), **a**_*i*_ is the corresponding column vector of direct genetic effects, and **e**_*i*_ is an uncorrelated vector of residual environmental effects; primes denote traits of interacting individuals of another species. The matrix **Φ**_*xy*_ quantifies the effect of traits in an interacting individual of species 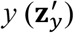 on the expression of traits in a focal individual of species *x* (Fig. 2, Table 2), while the matrix **Φ**_*yx*_ quantitifies such effects in the opposite direction. Thus, **Φ**_*ij*_ is an interspecific analog of the matrix of conspecific indirect genetic effects, **Ψ** (Moore et al. 1997). The individual elements of **Φ**_*ij*_, which we write as 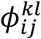, represent partial regression coefficients of trait *l* in species *j* on trait *k* in species *i*. When traits are standardized to the same scale, these coefficients will typically be limited to a range of −1 to 1. Although we focus on interactions between pairs of individuals for simplicity of notation, this model could be easily extended to describe interactions with multiple individuals (cf. McGlothlin et al. 2010). For more diffuse interactions (e.g., alewife and *Daphnia*), the elements of **Φ**_*ij*_ could represent a weighted average of effects from integrating across the phenotype distribution of the interacting population.

**Table 2.** Definition of key parameters and expressions.

| Expression / parameter | Biological definition |
| --- | --- |
| $\mathbf{z}_x, \mathbf{z}_y$ | Vectors of individual trait values for species $x$ and $y$, with a prime when present denoting partner traits. |
| $\mathbf{a}_x, \mathbf{a}_y$ | Vectors of individual breeding values for species $x$ and $y$. |
| $\Phi_{xy}, \Phi_{yx}$ | Interspecific indirect effects (in matrix form). The effect of traits in individuals of species $y$ on trait expression in species $x$, and vice-versa. |
| $\beta_{xx}, \beta_{yy}$ | Vectors of within-species natural selection gradients for each species. |
| $\beta_{xy}, \beta_{yx}$ | Vectors of cross-species selection gradients for each species. The effects of traits in species $y$ on fitness of individual species $x$, and vice-versa. |
| $\mathbf{G}_{xx}, \mathbf{G}_{yy}$ | Genetic covariance matrices of traits for species $x$ and $y$. |
| $\mathbf{G}_{xy}, \mathbf{G}_{yx}$ | Covariance matrices for breeding values of interacting individuals of species $x$ and $y$ within a population. |
| $\text{Cov}(\Delta\bar{z}_x, \Delta\bar{z}_y)$ | Coevolutionary covariance, defined as the covariance in evolutionary change between interacting species, across populations. When standardized by the variances in evolutionary response for each species, becomes a scale-free correlation. |
| $\text{Cov}(\beta_{xx}, \beta_{yy}),$ | Between-species, among-population covariance in selection gradients (shown here for within-species selection) |

Assuming that the mean residual environmental effect is zero, the population mean phenotype vector for each species is

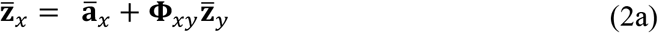

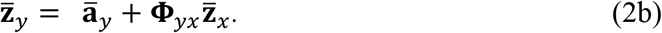

Noting the change in the mean additive genetic value in each species is 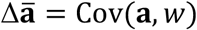 (Robertson 1966, Price 1970, 1972) where *w* is relative fitness, we can now define evolutionary change in the multivariate mean phenotype as

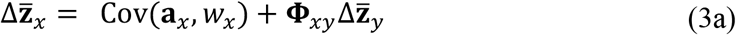

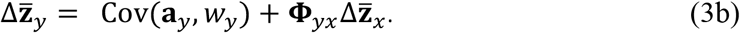

In equation (3), the first term on each right-hand side describes direct change due to natural selection on the focal species, and the second term describes indirect change due to the product of IIGEs and phenotypic change in the interacting species. This second term is analogous to transmission bias (Fisher and McAdam 2019), and in this case, specifically describes changes in one species that are induced by evolution in an interacting species. It can be seen from equation (3) that species will coevolve whenever there are IIGEs in both species, because the change in phenotypic mean in species *x* depends upon the change of mean in species *y* and vice versa whenever both **Φ**_*ij*_ ≠ **0**. Expanding equation (3) by substitution results in explicit equations for evolutionary change in each species,

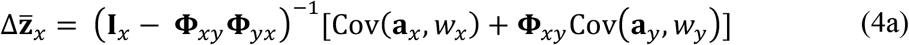

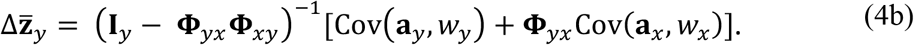

where **I**_*x*_ and **I**_*y*_ are identity matrices with dimensionality equal to the number of traits in the two species and the multiplier (**I** – **Φ**_*ij*_ **Φ**_*ij*_)^−1^ quantifies the feedback effect of reciprocal IIGEs. Equation 4 illustrates that total evolutionary change in a species is determined both by change in the breeding value of that species (Cov(**a**_*i*_, *w_i_*)) and by change in the breeding value of the interacting species, mediated by IIGEs (**Φ**_*ij*_Cov(**a**_*j*_, *w_j_*). Whenever IIGEs occur in both species such that **Φ**_ij_ **Φ**_*ji*_ ≠ 0, this multiplier alters the total amount of evolutionary change in both species. In order for such an effect to arise, there must be a feedback loop in phenotypic expression. The simplest of these arises when there are two traits with reciprocal IIGEs, such that trait *k* in species *i* affects trait *l* in species *j* and trait *l* in turn influences the expression of trait *k*. In general, when 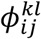 and 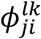 are of the same sign, the magnitude of evolutionary change will be enhanced, and when they are of opposite signs, evolutionary change will be diminished (Fig. 3).

To expand equation (4), we define the fitness of each interacting species using the linear equations

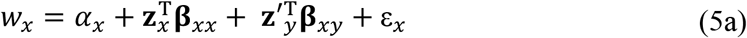

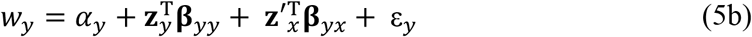

where *w* is individual relative fitness, *α* is an intercept, ε is an error term, and the superscript T denotes transposition. Directional selection in each species is partitioned into two selection gradients. First, the within-species selection gradients (**β**_*xx*_ and **β**_*yy*_) describe the direct effects of an individual’s traits on its own fitness (Fig. 2; Lande and Arnold 1983). The cross-species selection gradients (**β**_*xy*_ and **β**_*yx*_) relate the fitness of a focal individual of one species (*x* or *y*) to the traits of individuals of the other coevolving species (*y* or *x*) (Fig. 2). Both of these are linear components of selection; we consider nonlinear terms (interactions for fitness between focal and partner traits) below (see Incorporating Specific Fitness Models). The cross-species selection gradient is analogous to the directional social selection gradient in within-species models (Wolf et al. 1999) and if pairs or groups of interacting individuals can be identified in a natural population, can be estimated in a similar way (cf. Ridenhour 2005).

**Figure 3.**
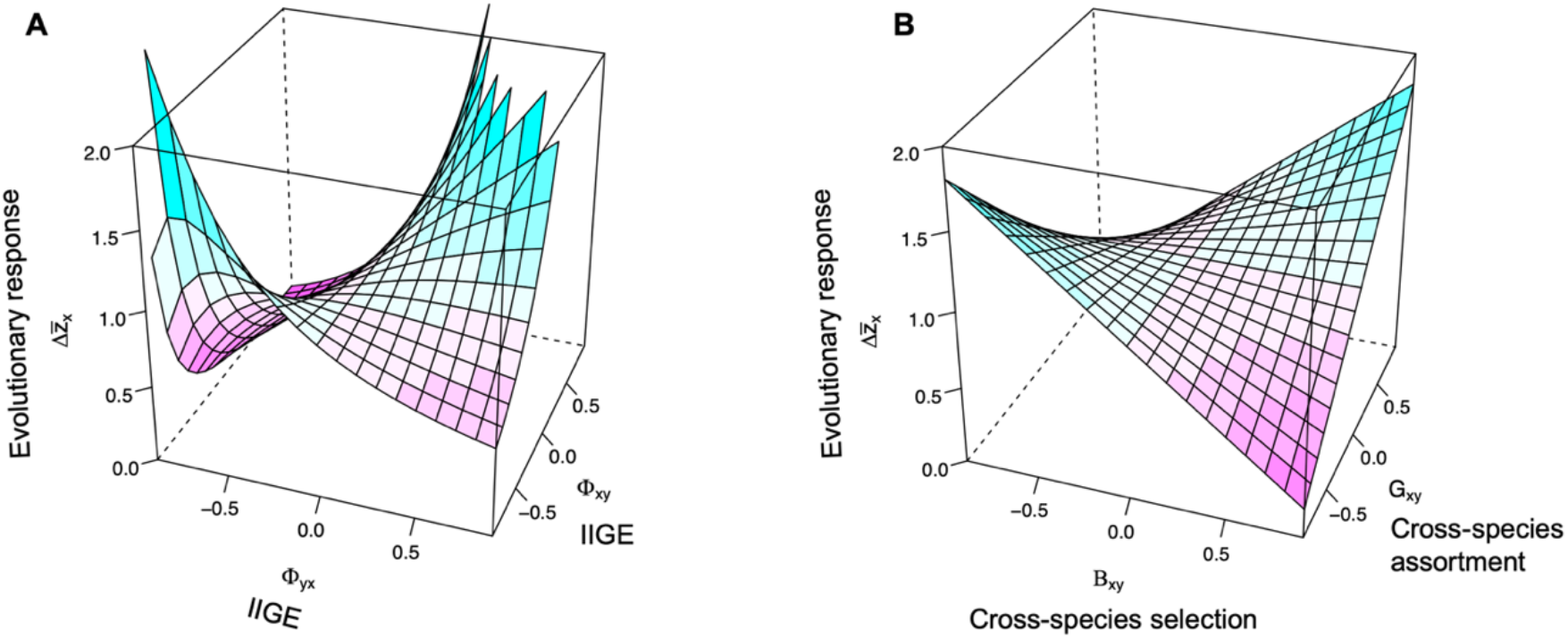
Reciprocal IIGEs and cross-species selection change evolutionary response in a single species. Panels show the separate effects on evolutionary response in species x of indirect genetic effects (Panel A) and cross-species selection with genetic assortment (Panel B). Panel A shows the effects of reciprocal IIGEs holding all other evolutionary parameters constant, and assuming no cross-species selection. Panel B shows the effects of cross-species selection imposed by species y on species x, in combination with the genetic assortment between interactants of the different species, and assuming IIGEs are absent. In both panels, *G_x_* = *G_y_* = 1, β_*yy*_ = β_*xx*_ = 1.

We now substitute our definitions of fitness into equation (4) and expand, yielding predictive equations for evolutionary change in each species:

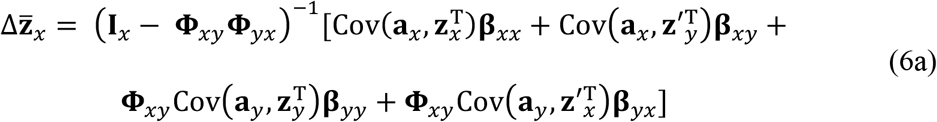

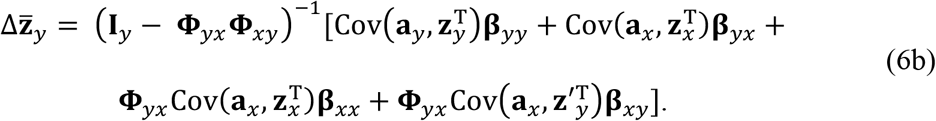

Each of these equations consists of four terms representing four components of the total response to selection. The first term 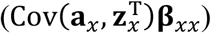 represents response to within-species selection (**β**_*xx*_ or **β**_*yy*_), the second 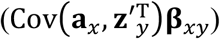 represents response to cross-species linear selection (**β**_*xy*_ or **β**_*yx*_), and the last two terms 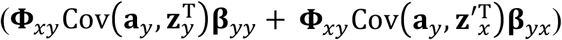 represent the component of change caused by the change in mean of the interacting species. Note that the third and fourth terms of the first part of equation (6a) equal the first two terms of (6b) multiplied by the IIGE coefficient **Φ**_*xy*_. Equation (6) also shows that the change in response to within-species selection depends on the covariance of additive genetic value with the phenotype of the same species, while change in response to cross-species selection depends on the covariance of additive genetic value with the phenotype of the opposite species.

To determine the components that give rise to these covariances, we expand our definition of the phenotypes in equation (2) and rearrange, yielding

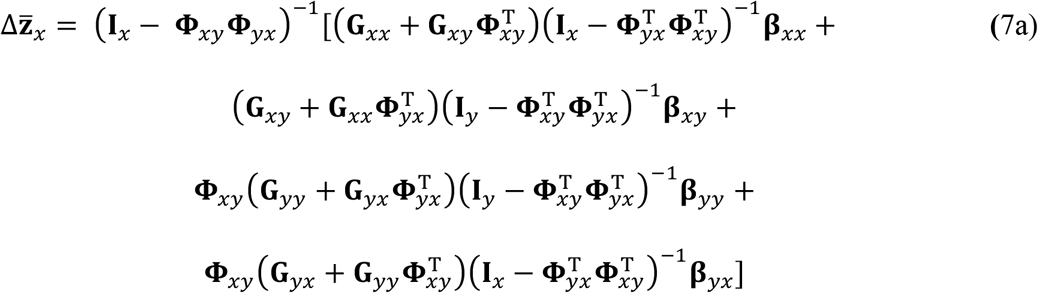

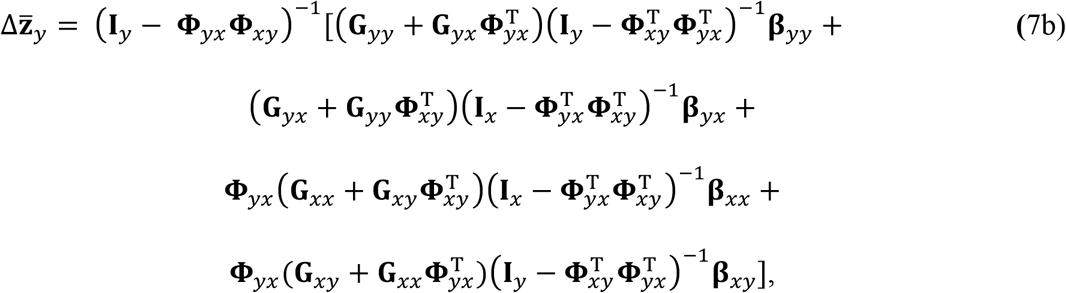

where **G**_*xx*_ and **G**_*yy*_ represent within-species genetic (co)variance matrices and **G**_*xy*_ and **G**_*yx*_ represent cross-species genetic covariance. Note that in a model of pairwise interactions, 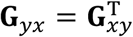.

The cross-species genetic covariance represents a covariance of breeding values between the species within the population of interest. This covariance may arise via any mechanism that leads to nonrandom genetic assortment between species *x* and *y* with respect to traits at the phenotypic interface of coevolution. A variety of phenomena could lead to such assortment including behavioral preference for certain traits in heterospecific partners, fine-scale population structure, habitat preference, and vertical transmission of symbionts. We elaborate on the contribution of **G**_*xy*_ and **Φ** to phenotypic assortment between individuals of interacting species (*C_xy_*) in equation (A1). However, there is currently limited evidence of this type of interspecific genetic assortment, as few investigators seemed to have attempted to measure such a covariance. We therefore investigate dynamics in both the presence and absence of such assortment in our model development below.

Response to within-species selection depends on the sum of within-species genetic variance, **G**_*yy*_ or **G**_*xx*_, and 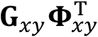 or 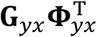, which represent an interaction between IIGEs and cross-species genetic covariance. Response to cross-species selection depends on the sum of cross-species genetic variance and 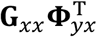 or 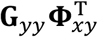, which describes the genetic variance created by IIGEs. Each term in equation (7) also contains an additional feedback multiplier, 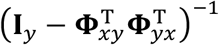, which further enhances the response to selection when there are reciprocal IIGEs. Because of this additional multiplier, when IIGEs are reciprocal and of the same sign, their effects on can be massive, mirroring the effects observed for within-species IGEs (Moore et al. 1997, McGlothlin et al. 2010). As in previous equations, the last two terms in equation (7) represent a sort of evolutionary feedback that occurs across generations and is only present when there are IIGEs. These effects of **Φ** and cross-species selection on evolutionary response are illustrated in Fig. 3.

### Correlated evolution between interacting species: the coevolutionary covariance

A key feature of coevolving lineages is correlated evolution across populations subject to varying ecological conditions. Here, we seek to understand how the equations of selection response can be used to understand how selection and IIGEs contribute to this shared among-population divergence. We can explore the contribution of interspecific social effects to correlated evolution between interacting species by solving for the covariance in evolutionary change between interacting species, 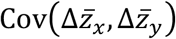, which we call the coevolutionary covariance (with the caveat that this covariance also reflects changes due in part to effects of IIGEs). Such a covariance represents the expected pattern of reciprocal phenotypic change through time in a single pair of populations of two species experiencing fluctuating selection, or perhaps more importantly, across a set of populations in space under varying selection pressures. Although mathematically equivalent, we focus on the latter scenario for its relevance to understanding geographic variation among populations.

The coevolutionary covariance reflects the degree to which divergence among population means in two species has occurred jointly. High absolute values of the coevolutionary covariance indicate tightly-coupled coevolutionary change between the two species, whereas values around zero indicate that evolutionary change occurs independently. When scaled by the total amount of population divergence (variances in 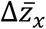 and 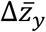), this becomes a scale-free correlation describing the proportion of total divergence shared between the two species. Importantly, we expect the coevolutionary covariance to be often be related to the covariance in population means, 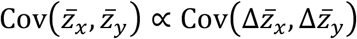 because, for example, 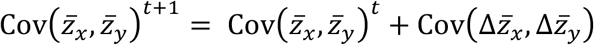 (under the assumption 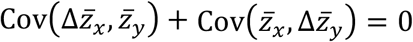). Thus, expanding 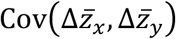 allows the possibility to assess how selection and IIGEs contribute to patterns of geographic variation in species mean phenotypes.

To simplify our analysis, we use models of a single trait in each species. Developing a full equation including all sources of covariance quickly becomes cumbersome, so we focus on three instructive special cases that illustrate the explicit impacts of considering crossspecies selection and interspecific interacting phenotypes. The simplest case occurs when there are no IIGEs (*ϕ_xy_* = *ϕ_yx_* = 0) and heterospecific interactions occur at random within populations (*G_xy_* = *G_yx_* = 0). If we make the simplifying assumption that genetic variance does not differ among populations, the only source of covariance in selection response is covariance in direct within-species selection, or

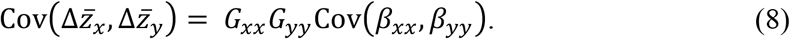

Equation (8) represents what is usually thought of as one source of correlated evolution between interacting species: predictable variation in the type of selection occurring across space. Empirical data suggest that selection varies substantially in magnitude across space (Siepielski et al. 2013) and that spatially autocorrelated biotic selection plays a substantial role in driving divergence in trait means (Urban et al. 2011). Importantly, not just any variance in selection will do. In order to create covariance in evolutionary change, selection must covary between the interacting species. Cross-species selection, however, does not play a role in equation (8) because it does not contribute to an evolutionary response to selection in the absence of IIGEs and cross-species genetic covariance. Equation (8) is analogous to the results of Zeng’s (1988) model of correlated trait evolution under directional selection, although for the case of traits expressed in different species.

To see an effect of cross-species selection, we first add cross-species genetic covariance but no IIGEs. For a one-trait (per species) model, *G_xy_* = *G_yx_*. Adding this effect leads to three new sources of covariance among populations:

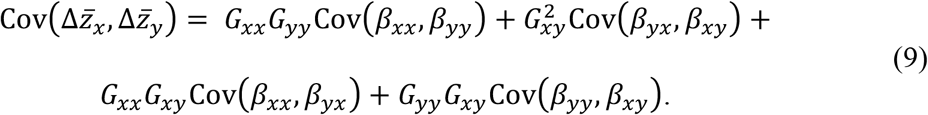

The second term in equation (9) 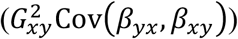 shows that cross-species selection will contribute to the coevolutionary covariance only when there is cross-species genetic covariance for the traits that affect fitness. Such covariances occur if, for example, parasite and host genotypes associate nonrandomly for the traits that influence their partner’s fitness. The third and fourth terms (*G_xx_G_xy_*Cov(*β_xx_,β_yx_*) + (*G_yy_G_xy_*Cov(*β_yy_,β_xy_*)) represent a relationship between the effect of species on itself and on its heterospecific partner. These will be nonzero if populations with strong within-species selection also exhibit strong crossspecies selection.

The most interesting effects on the coevolutionary covariance occur when we add IIGEs (*ϕ_xy_* and *ϕ_yx_*). With *G_xy_* = 0, an appropriate simplifying assumption given that the prevalence of *G_xy_* is uncertain and is likely to be transient through time, the covariance among populations in evolutionary response to selection is

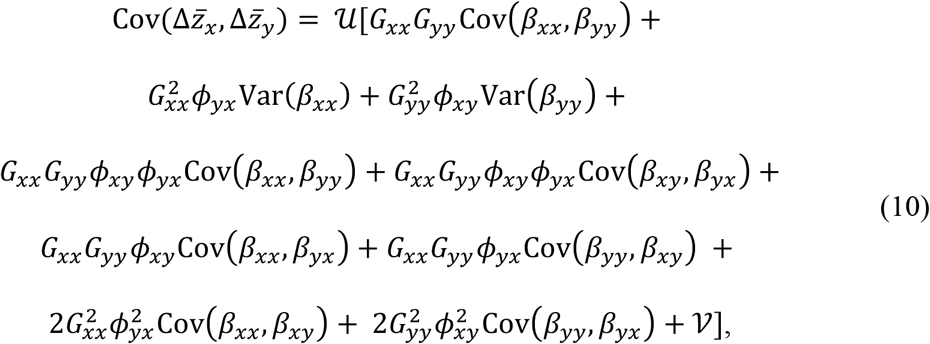

where feedback effects of IIGEs are described by 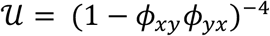 and 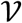 collects negligible third- and fourth-order *ϕ* terms:

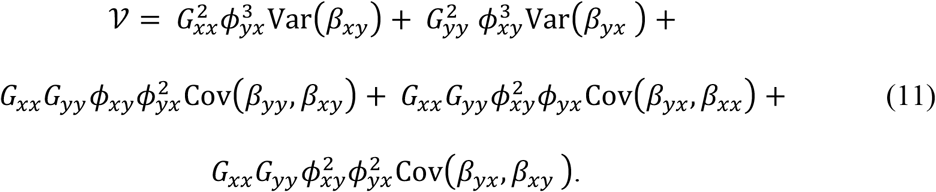

The first term in equation (10) is identical to equation (8) and represents covariance in direct within-species selection. The second 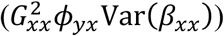 and third 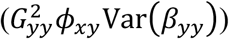 terms show that in the presence of IIGEs, simple variance in within-species selection across populations can generate a coevolutionary covariance (or correlation, Fig. 4A; corresponding evolutionary rates and covariances are plotted in Figs. S1 and S2, respectively). This covariance in evolutionary response occurs as a necessary consequence of the dependence of the trait mean of one species on that of the other. Thus, in the presence of IIGEs, a coevolutionary covariance may occur even when selection is uncorrelated between the species, and in extreme cases, when only one species varies in selection or when only one species has genetic variation (Fig. 4B).

**Figure 4.**
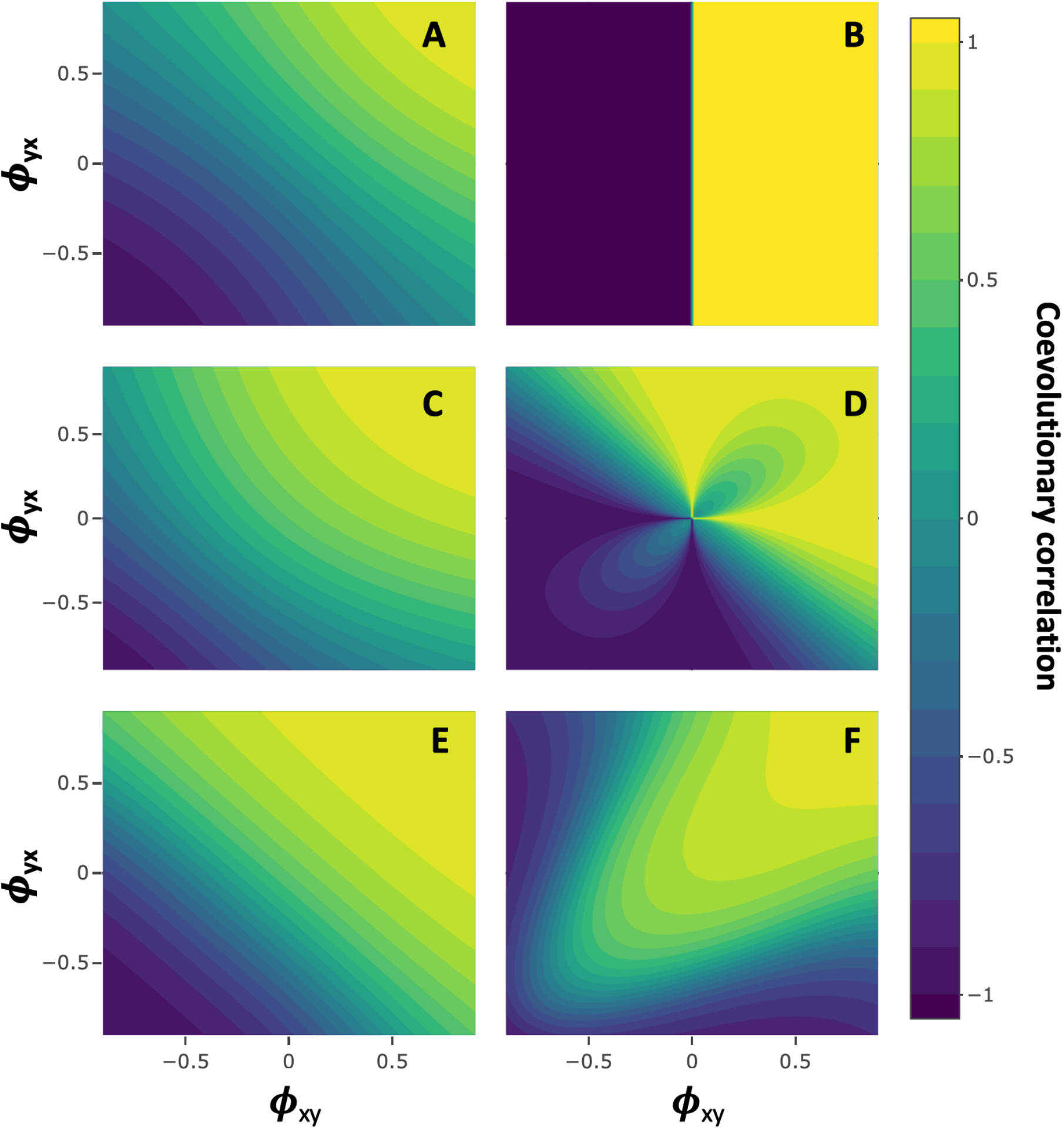
IIGEs drive and mediate coevolution between interacting species. Panels show the coevolutionary correlation between two interacting species as a function of the indirect genetic effect parameters *ϕ*, from equation (10) standardized by the evolutionary rates. In the absence of social interaction effects, correlated evolution (coevolution) between species is driven entirely by covariance in natural selection between the species (see text). Reciprocal IIGEs can generate coevolution even when there is not covariance in natural selection (Panel A), and can even drive coevolution even when one species lacks genetic variance (Panel B). IIGEs modify observed coevolutionary patterns when natural selection does covary (Cov(β_*xx*_, β_*yy*_) = 0.5; Panel C). When cross-species selection and IIGEs act together, coevolutionary patterns are a complex third order polynomial (Panel D; Cov(β_*xy*_, β_*yx*_) = 0.5, Var(β_*xy*_) = Var(β_*yx*_) = 1). When natural and cross-species selection both act (Var(β_*xy*_) = Var(β_yx_) = Var(β_*yy*_) = Var(β_*xx*_) = 1) and covary positively (0.5), (Panel E), effects of IIGEs become stronger in comparison to the case of (co)variance in natural selection alone. When covariance between natural and cross-species selection is negative (−0.8) a ridge is observed (Panel F).

The fourth and fifth terms (*G_xx_G_yy_ϕ_xy_ϕ_yx_*Cov(*β_xx_,β_yy_*) + (*G_xx_G_yy_ϕ_xy_ϕ_yx_*Cov(*β_xy_,β_yx_*)) show the effects of reciprocal IIGEs. When traits in the two species exist in a feedback loop, the effect of covariance in within-species selection is amplified if *ϕ_xy_* and *ϕ_yx_* are of the same sign and diminished if they are of opposite signs. The combined effects of the second, third, and fourth terms lead to a complex relationship between IIGEs and the coevolutionary covariance when within-species selection varies (Fig. 4C). In some cases, IIGEs can even reverse the sign of the correlation that would be expected in their absence (Fig. 4C). In addition, reciprocal IIGEs may cause covariance in cross-species selection to contribute to the coevolutionary covariance, causing an even more complex relationship (Fig. 4D).

The last four terms in equation (10) show that in the presence of IIGEs, covariance between within-species and cross-species selection may contribute to the coevolutionary covariance (Fig. 4E-F). Because IIGEs inherently tie together the evolutionary responses of the two species through their effects on the coevolutionary covariance, this can occur both when there is covariance in gradients across species (terms 6 and 7) and when there is covariance in gradients in the same species (terms 8 and 9). The total effect of IIGEs may be quite complex when cross-species selection varies, with subtle changes in *ϕ* resulting in dramatic changes in the expected among population correlation (Fig. 4D, E-F).

An equation incorporating both IIGEs and *G_xy_* quickly becomes unwieldy, but we present a compact form containing 10 (co)variance terms as equation (A2), which illustrates that when present, non-random assortment and IIGEs together interact in complex ways to influence coevolution.

### Incorporating specific fitness models

Coevolutionary models often posit complex relationships between interacting phenotypes and fitness (Nuismer 2017). Although the selection model we present here is linear, more complex relationships can be incorporated by translating specific fitness functions to selection gradients. The adaptive landscape represents the theoretical relationship between a population’s mean fitness and its phenotypic mean (Arnold et al. 2001). Selection gradients represent the partial slope of the adaptive landscape with respect to a given phenotype, and thus the multivariate selection gradient may be calculated using a vector of partial derivatives if the adaptive landscape can be written as a differentiable function (Lande 1979, Lande and Arnold 1983). In many cases, the multivariate selection gradient may be calculated using partial derivatives of the individual fitness function as well (Lande and Arnold 1983, Abrams et al. 1993, McGlothlin et al. 2021), evaluated over the phenotypic distribution (Phillips and Arnold 1989). Once selection gradients have been calculated for a given model, they may be substituted into equations (6–7) to explore the effects of a given fitness function on selection response and the coevolutionary covariance (see also Brodie and Ridenhour 2003). This exercise also allows us to explore the effects of adding IIGEs to an existing coevolutionary model.

First, we consider a “phenotypic difference” model of ecological trait interaction (Nuismer et al. 2007, Nuismer 2017), where absolute fitness is a function of the difference between interacting traits, *W_x_* ∝ exp (z_*x*_ – z_*y*_) and *W_y_* ∝ exp(z_*y*_ – z_*x*_). Using the logarithm of the fitness function to calculate relative fitness (Lande and Arnold 1983),

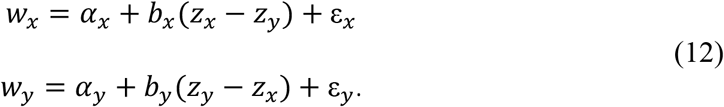

where *b_x_* and *b_y_* are constants that may vary across populations. Differentiating, the selection gradients are then *β_xx_* = −*β_xy_* = *b_x_* and *β_yy_* = −*β_yx_* = *b_y_*. Substituting these selection gradients into equation (7) is trivial. However, is worth noting that under this fitness model, the coevolutionary covariance simplifies to a function of just three (co)variance components,

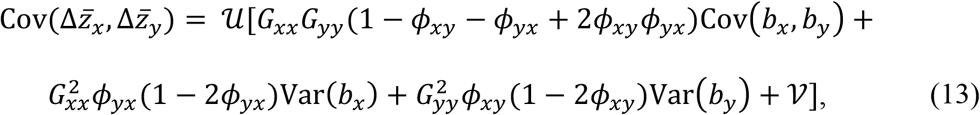

which corresponds to a relationship with *ϕ* as in Figs. 4F, S1F, and S2F. This analysis illustrates that reciprocal IIGEs have greatest impact on mediating coevolutionary outcomes in a trait-matching models when IIGEs are similar in both sign and magnitude. Biologically, such a situation corresponds to a scenario where, for example, trait expression is reciprocally escalated in response to heterospecific partners.

Another important class of fitness models to consider is the case of nonlinear fitness interactions. Nonlinearity is fundamental to many coevolutionary models that invoke epistatic interactions across species’ genomes, such as the trait matching model for quantitative traits, or the single-locus matching-allele and gene-for-gene models of hostparasite coevolution (Dybdahl et al. 2014). For example, in the traditional model of trait matching, absolute fitness is assumed to be *W_x_* ∝ exp (−(z_*x*_ – z_*y*_)^2^) and so relative fitness can be described by

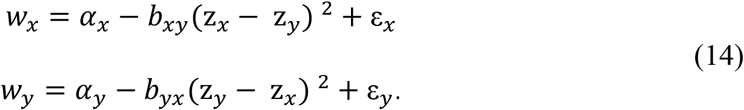

In this model the sign of the *b* terms correspond to different types of biological interaction; for example, in a mutualism, both species may take positive values of *b* and thus have a fitness peak when their trait value matches with their heterospecific partner. Taking partial derivatives over the phenotypic distribution yields the selection gradients

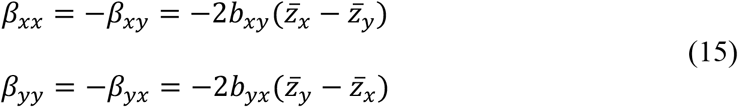

In the simplest case of no IIGEs, *G_xy_* = 0, and no variation in *b*, these gradients yield

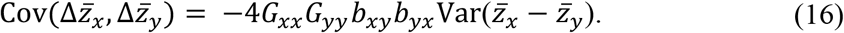

Thus, within the limits of genetic variation and ignoring stochastic forces (as we have done), evolutionary response will be perfectly correlated in the phenotypic matching model. When *b_xy_* and *b_yx_* share the same sign, this covariance in evolutionary response will be negative because each species will be evolving towards (or away) from each other in trait space. This covariance in evolutionary response occurs even without variation in *b* or covariance in trait means. These results are broadly consistent with other analyses of the trait matching model, which indicate that this model can generate relatively strong covariance in phenotypic means across populations (Nuismer et al. 2010). Adding indirect genetic effects and assuming *G_xy_* = 0 and noting that 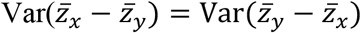,

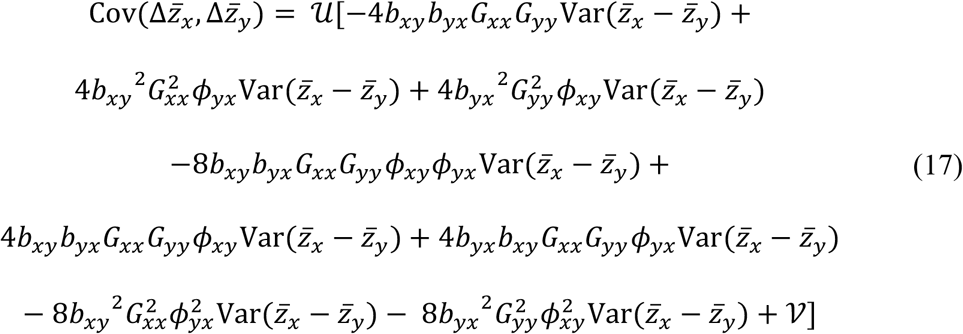

This expression again shows the covariance in evolutionary response occurs even in the absence of (co)variance in *b*, and the addition of IIGEs alter the magnitude of the covariance substantially. Analysis of more complex cases of the trait matching model, for example where selection varies across space and/or across terms in the expanded polynomial (z_*x*_ – z_*y*_)^2^, would be straightforward in this framework.

As a final example of how this approach can be used to understand simple variations on classic coevolutionary models, we focus on a multiplicative model of trait interaction, where fitness depends on the interaction between traits of the two species. Consider a fitness model where relative fitness depends solely on the product of the two phenotypes:

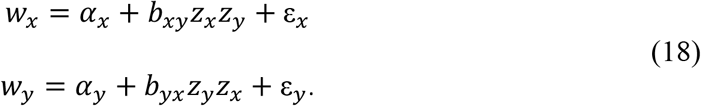

This model is conceptually and mathematically similar to the previous trait matching model, but importantly lacks stabilizing selection terms (e.g., 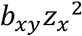) present in the trait matching model. Because of this, directional natural selection depends only on the means of the other species:

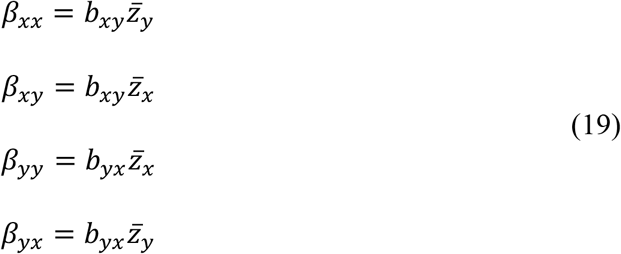

This relationship may cause selection to (co)vary across populations even when *b_xy_* and *b_yx_* are homogeneous. In the absence of IIGEs, this simplest case would lead to a coevolutionary covariance defined by

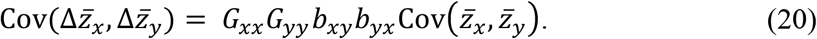

Thus, any initial covariance in the population means leads to a covariance in the response to selection across species, lending a runaway aspect to the coevolutionary covariance. Adding IIGEs, this becomes

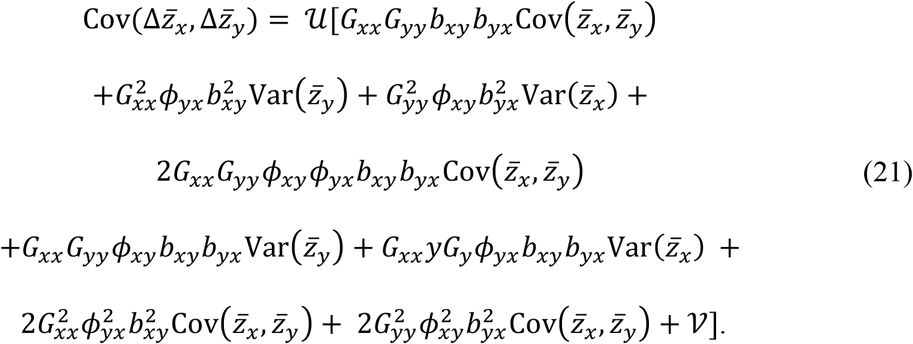

This equation again shows that when IIGEs are present, any variance across populations in the trait mean of either species leads to a cross-species covariance in the response to selection (terms 2, 3, 5, and 6). Cross species covariance in selection response is also mediated further by covariance in trait means in this model (terms 1, 4, 7, and 8) as well as higher order products of IIGEs captured in *V* (equation 11). These effects may lead trait means to become correlated across populations in future generations. This analysis, particularly contrasting equations 16 vs. 20 and 17 vs. 21, also reveals that coevolutionary dynamics can be substantially different even across models that share a common polynomial relative fitness function.

## Discussion

Our model adapts the theory of trait-based intraspecific social evolution to the phenotypic interface between two coevolving species. We show that two forms of interspecific interaction, interspecific indirect genetic effects (IIGEs) and cross-species selection (analogous to within-species social selection), both contribute to correlated evolution between interacting species. Our analysis shows that reciprocal IIGEs modulate selection response, suggesting that IIGEs may play a major role in generating and mediating patterns of correlated evolution between species. Further, we show that constant (across space) IIGEs can generate a coevolutionary covariance even in the absence of covariance in selection, or even the absence of genetic variance in one species. When selection does covary between species across populations, reciprocal IIGEs will promote changes in the magnitude of coevolution and even reversals in the expected among-population covariance. IIGEs also allow cross-species selection, which we model as the effect of the traits of one species on the fitness of another, to influence evolutionary response. Such a response may also be mediated by cross-species genetic assortment between interacting individuals. When IIGEs and crossspecies selection act together, effects on the coevolutionary covariance can be complex, with dramatic changes in the expected sign and magnitude of correlated evolution occurring with subtle changes in these parameters of interspecific social interaction. Our results indicate that whenever coevolving species socially interact to modify expression of one another’s phenotypes, these interspecific social interactions are key to understanding coevolution.

IIGEs represent a scenario where phenotypes in individuals of one species influence trait expression of individuals of another species. Thus, IIGEs are a specific type of environmental effect (Moore et al. 1997, Drown and Wade 2014) where the environment is the phenotypic value of the interspecific individual(s) with which an organism interacts. To our knowledge, this type of environmental effect on between-species coevolution has been considered in only two other theoretical studies (Scheiner et al. 2015, O’Brien et al. 2021; but see Shuster et al. 2006, Witham et al. 2020 for a variance-partitioning approach). Scheiner et al. (2015) consider a special case of our model, where evolvable IIGEs are present in only one of the two interacting species. In this non-reciprocal model, they show a much more limited role for IIGEs in coevolution. Our results are broadly consistent with this conclusion, in that IIGEs in only a single species do not generate the reciprocal effects that lead to massive inflation of evolutionary response. However, IIGEs in only a single species (e.g., **Φ**_*xy*_ = 0, **Φ**_*yx*_ ≠ 0) still play a role in mediating response to interspecific social selection whenever interspecific social selection is a function of individual trait values (as opposed to the population mean, as modeled by Scheiner et al. 2015). We also note that our fully multivariate model accommodates the possibility that reciprocal IIGEs act across different types or numbers of traits in the two interacting species. More recently, O’Brien et al. (2021; see also Queller 2014) developed a model of coevolution between host plant and microbial symbionts. Their parameterization differed from ours in that they consider evolution of a single joint trait governed by genetic variation in host and symbiont, and so is most directly applicable to plant-microbe systems or other intimate interactions. Nonetheless, their model shows an important role for reciprocal fitness feedbacks, consistent with the conclusions of our trait-based model.

Within-species models of interacting phenotypes clarify the line between genetic and environmental effects, furthering an understanding of how genotypes expressed in an individual that act as environments for other individuals can influence phenotypic change (Wolf et al. 1998, Wolf 2003). That is, IGEs are genetically-based environments that influence phenotypic expression during interactions. Conceptually, this relationship is similar to genetically based plasticity. Extending these types of models to the case of interspecific interaction carries similar challenges and benefits. We have referred to reciprocal change in phenotypic means between interacting species as “coevolution,” even when these effects are mediated by IIGEs. This is a broad use of the term coevolution, as IIGEs are an environmental effect that can themselves evolve, again similar in concept to phenotypic plasticity. However, a critical difference is that changes in phenotypic response mediated by IIGEs represent changes driven by evolution of an interacting species (made clear in equation 4). Our point is not to broaden the definition of coevolution, but rather to highlight that IIGEs can have substantial impact on patterns of phenotypic divergence in coevolving species. For example, our model highlights that genetic divergence across populations of a single species is sufficient to generate tightly-coupled patterns of correlated change in an interacting species, a result that suggests the challenges of interpreting correlated phenotypes as evidence of genetic response to reciprocal selection may be even greater than already appreciated (e.g. Nuismer et al. 2010, Gomulkiewicz et al 2007, Janzen 1980).

Interspecific indirect genetic effects, or at least the potential for a prevalence of such effects, appear to be commonplace in many biological systems. In Table 1, we provide in a breakdown of types of biological interaction in which there is a large literature suggesting importance of IIGE-like phenomena. These types of effects on trait expression across species, widely appreciated in their own specific contexts (Weis and Abrahamson 1986, Peacor and Werner 2001, Werner and Peacor 2003, Chen 2008, Thomas et al. 2012, O’Brien et al. 2021), have taken on a variety of different forms. We suggest that these disparate biological phenomena may nonetheless share a commonality—reciprocal effects on trait expression across interspecific partners—that we have shown can affect the coevolutionary process in dramatic, and in some cases predictable, ways.

Cross-species selection features prominently in verbal descriptions of the coevolutionary process (Thompson 1982), and we show that such selection is especially important in the presence of interspecific indirect genetic effects. When individual trait values of one species affect individual fitness of another, focal species, this cross-species selection can manifest evolutionary change in the focal species when there is phenotypic assortment between interspecific interactants. This assortment, analogous to that required for evolutionary response to social selection within species (Wolf et al. 1999, McGlothlin et al. 2010, Brodie et al. submitted), can be generated directly by a non-random genetic assortment, or via IIGEs. Examples of processes that could generate direct genetic assortment between interacting individuals of two different species include shared genetic structure, to the extent that such structure manifests assortment of breeding values for the relevant traits. Such shared genetic structure could arise through shared features of the environment that limit gene flow and panmictic mating in both species, or alternatively, through variation in habitat preference across individuals of both species. Direct genetic assortment could also arise through behavioral preference for certain trait values in heterospecific partners. Such preferences may be especially common in predator-prey interactions, where, for example, predator body size may be expected to coevolve with behavioral preference for prey size (Troost et al. 2008). Currently, it is unclear how common this type of cross-species genetic assortment may be, although in part this likely reflects a lack of studies that have attempted to measure assortment between breeding values of individuals of separate species. Moreover, when it does occur, such assortment is likely to be transient because it does not rely on transmission of pleiotropic alleles that may stabilize within-species genetic correlations over multiple generations. The substantial evidence for IIGEs, but limited evidence of *G_xy_* suggests that IIGEs may play a prominent role in mediating any realized response to crossspecies selection.

Our results also indicate that nonlinear effects on cross-species selection can contribute to coevolution even in the absence of genetic assortment or IIGEs. This form of selection corresponds to an interaction between focal and interspecific-partner trait values for focal individual fitness. The effect of this form of cross-species selection on evolutionary response in a population depends on the mean genotype of the other species, and thus represents a diffuse effect of population mean phenotype of the coevolving species. Such interspecific interactions are potentially less intimate, for example diffuse predator chemical cues in aquatic environments, than the individual level interactions (e.g., of host and parasite) required to generate response from linear cross-species selection. Across populations, nonlinear cross-species selection contributes to coevolution via covariance in mean genetic values and/or linear selection between the species. This result is consistent with past models of coevolution, verbal and mathematical, that indicate trait interactions for fitness are a key feature of coevolution (Thompson 1982, 1994, 2005, Nuismer 2017), and in our model, such interactions lead to a dependence between selection in one species and the mean trait value of another. By defining these interaction terms in the framework of social evolution, our model adds to past work by indicating that reciprocal IIGEs can substantially increase the degree to which trait interactions for fitness contribute to reciprocal evolutionary change.

A key feature of our model is the development of a formal expression for the expected covariance in evolutionary response, 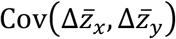. This coevolutionary covariance is expected to be a key contributor to generating among-population covariation in species mean trait values, a major focus in many empirical (Thompson 1994, 1995, Zangerl and Berenbaum 2003, Toju and Sota 2005, Hanifin et al. 2008, Hague et al. 2020) and theoretical (Nuismer et al. 2010, Nuismer and Week 2019, Week and Nuismer 2019) studies of coevolution. Importantly, similar to existing within-species models of among-population quantitative genetic variation (Zeng 1988, Chenoweth et al. 2010), defining this coevolutionary covariance illustrates how selection, IIGEs, and genetics may contribute to patterns of trait variation across populations.

Our model subsumes mechanistic detail into broad statistical descriptions of species interactions and thus provides a general description of how IIGEs and cross-species selection, when present, contribute to reciprocal evolutionary change and correlated evolution across populations. In contrast to our approach, some models of coevolution have focused instead on specific ecological mechanisms that generate trait-fitness relationships between interacting species (reviewed in Nuismer 2017). By highlighting the key parameters that contribute to coevolution—covariance in natural selection, covariance in cross-species selection, and IIGEs—our model indicates various pathways through which specific ecological mechanisms may affect coevolution. Our framework can be tailored to specific scenarios by substituting different fitness models into the general equations we present here.

Our model generates quantitative predictions for the shape of coevolution that are directly testable with empirical data because it focuses on estimable statistical effects of underlying ecological mechanisms rather than the mechanisms themselves, which are often unknown (Wade and Kalisz 1990). For example, using an empirical estimate of **Φ** (which could be measured using methods analogous to those used to measure within-species indirect genetic effects; Bleakley and Brodie 2009, McGlothlin and Brodie 2009), one could use matrix comparison of covariances among population means and the covariance terms presented here to quantitatively test the contribution of IIGEs to among-population covariance in selection response between two interacting species (e.g., see Chenoweth et al. 2010 for a within-species test of the predictions of Zeng’s 1988 model). More generally, Week and Nuismer (2019; see also Nuismer and Week 2019) have shown how datasets of among-population variation in trait means can be used to test for conformation to expectations from coevolutionary models. Concomitantly, our models show how environmental effects can be partitioned into terms describing genotypes of other species in the ecological community, which could be useful in understanding when and why evolutionary response fails to conform to predictions arising from the standard breeder’s equation.

Social interactions between individuals of the same species play a central role in the evolutionary process. Within a single lineage, indirect genetic effects and social selection fundamentally change selection response, the expression of genetic variance, and together determine the course of social evolution (Moore et al. 1997, Wolf et al. 1998, Wolf et al. 1999, McGlothlin et al. 2010). We have shown that these effects of interactions among individuals may transcend species boundaries and profoundly impact the dynamics of coevolution between interacting lineages.

## Author Contributions

All authors contributed to all aspects of the manuscript.

## Acknowledgements

Funding was provided by grants from the Royal Swedish Academy of Sciences and Swedish Research Council to S. De Lisle (VR registration number 201903706), the University of Connecticut and the NIAID (1R01AI123659-01A1) to D. Bolnick, and the National Science Foundation (DEB 1457463) to J. McGlothlin.

## Data Accessibility

No data to be archived

## Supplemental Figures

**Figure S1.**
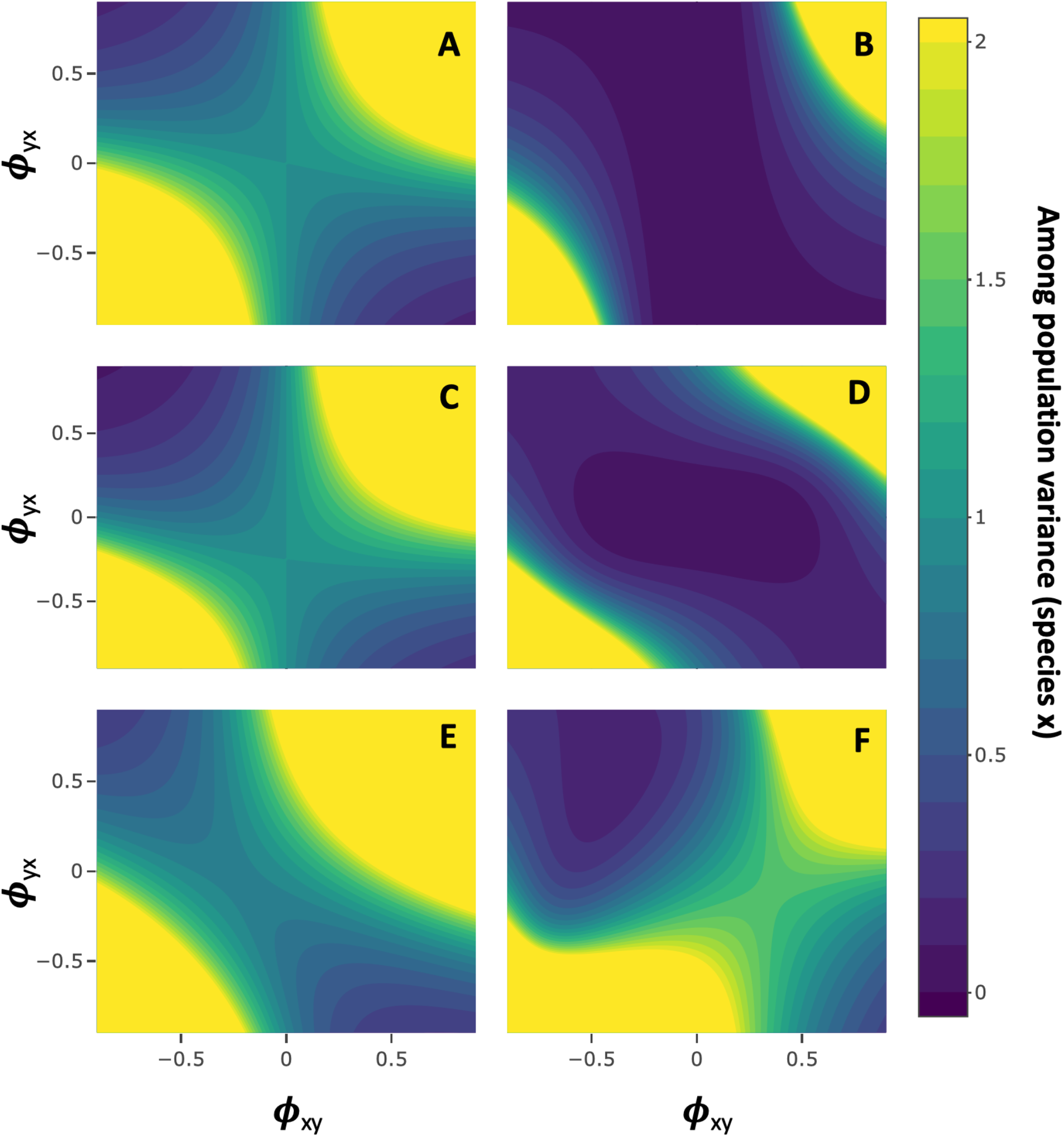
IIGEs accelerate evolutionary rate in a single species. Panels show the evolutionary rate, 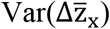, of species *x* as a function of the indirect genetic effect parameters *ϕ*, under the same parameter values as in Figure 3. Reciprocal IIGEs between interacting species generally accelerate evolutionary rate. Note that in the absence of any other effects, the evolutionary rate is equal to the variance in natural selection, which is unity in Panels A, C, E, and F. In panel D, evolutionary rate is driven entirely by cross-species selection and IIGEs. In panel B, where G_*xx*_ = 0, evolutionary rate in species *x* is driven entirely by reciprocal IIGEs and evolutionary change in species *y*.

**Figure S2.**
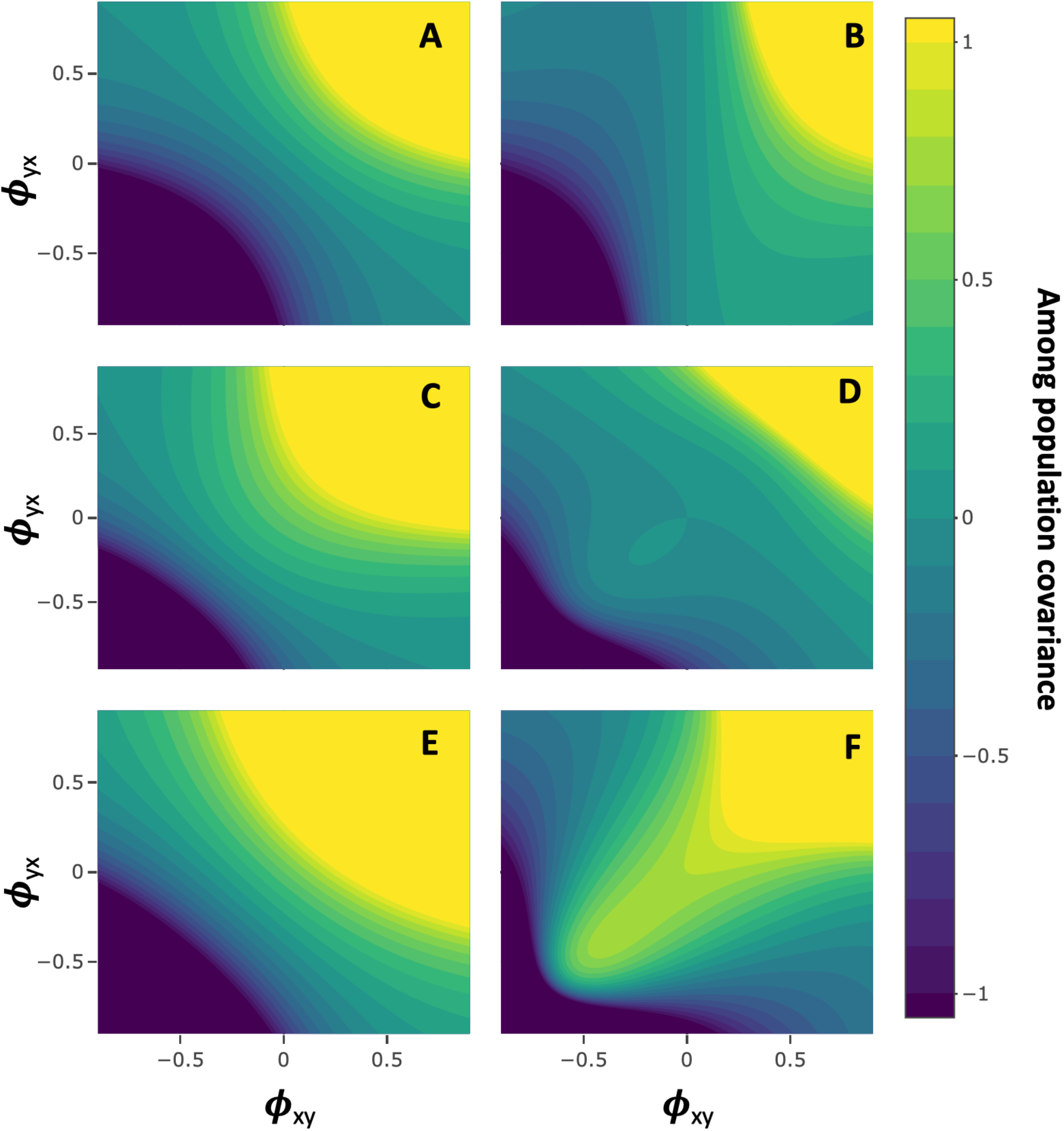
Among population covariances. Panels show the coevolutionary covariance between two interacting species as a function of the indirect genetic effect parameters *ϕ*, from equation (7) unstandardized. Parameter values are as in Figs. 4 and S1. For all panels, genetic variances were set to unity.

## Appendix

## A1. Phenotypic covariance between interacting species

For a single trait, we can solve for the covariance between *z_y_* and *z_x_* to partition the phenotypic covariance between individuals of two interacting species into terms describing the contribution of IIGEs and terms describing non-random genetic assortment. Assuming cross-species environmental covariance is zero,

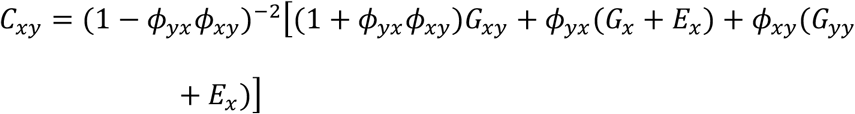

where *E_x_* and *E_y_* represent within-species environmental variance and *G_xy_* = *G_yx_*. When IIGEs are absent, non-random genetic assortment *G_xy_* is the sole contributor to the phenotypic association between individuals of coevolving species. When IIGEs are present, they can substantially change this phenotypic association.

## A2. Covariance in selection response with nonzero IIGEs and genetic assortment

We expand the covariance in evolutionary response in two species when IIGEs are present and constant and when genetic assortment *G_xy_* = *G_yx_* is present and constant,

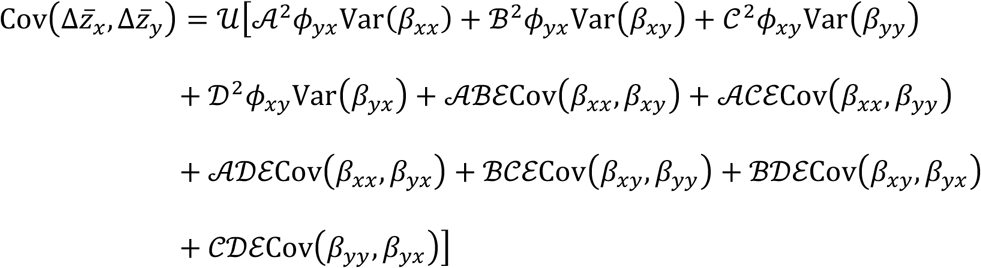

where

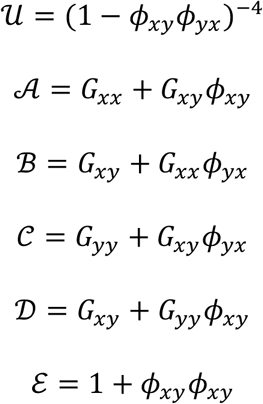

